# Distinguishing contemporary hybridization from past introgression with post-genomic ancestry-informative SNPs in strongly differentiated *Ciona* species

**DOI:** 10.1101/030346

**Authors:** Sarah Bouchemousse, Cathy Liautard-Haag, Nicolas Bierne, Frédérique Viard

**Author notes:** Correspondence : UMR7144, Equipe Diversité et Connectivité dans le paysage marin Côtier(DIVCO), CNRS-UPMC, Station Biologique de Roscoff, Place Georges Teissier, 29680 Roscoff, France.

## Abstract

Biological introductions bring into contact species that can still hybridize. The evolutionary outcomes of such secondary contacts may be diverse (e.g. adaptive introgression from or into the introduced species) but are not yet well examined in the wild. The recent secondary contact between the non-native sea squirt *Ciona robusta* (formerly known as *C. intestinalis* type A) and its native congener *C. intestinalis* (formerly known as *C. intestinalis* type B), in the western English Channel, provides an excellent case study to examine. To examine contemporary hybridization between the two species, we developed a panel of 310 ancestry-informative SNPs from a population transcriptomic study. Hybridization rates were examined on 449 individuals sampled in 8 sites from the sympatric range and 5 sites from allopatric ranges. The results clearly showed an almost complete absence of contemporary hybridization between the two species in syntopic localities, with only one first generation hybrid and no other genotype compatible with recent backcrosses. Despite the almost lack of contemporary hybridization, shared polymorphisms were observed in sympatric and allopatric populations of both species. Furthermore, one allopatric population from SE Pacific exhibited a higher rate of shared polymorphisms compared to all other *C. robusta* populations. Altogether, these results indicate that the observed level of shared polymorphism is more probably the outcome of ancient gene flow spread afterwards at a worldwide scale. They also emphasise efficient reproductive barriers preventing hybridization between introduced and native species, which suggests hybridization should not impede too much the expansion and the establishment of the non-native species in its introduction range.

## Introduction

Speciation is a gradual spatio-temporal process during which geographical or ecological isolation decrease gene flow between groups of individuals (Abbott *et al.* 2013). Species range shifts can deeply modify the evolution of these emerging species by promoting the formation of contact zones (Hewitt 2004; Maggs *et al.* 2008; Swenson & Howard 2005). In cases of species that are not fully reproductively isolated, interspecific gene flow occurs across hybrid zones (Barton 1979; Hewitt 2011). Hybridization and introgression processes between species in contact zones are particularly interesting to provide insights about the relative role of intrinsic and extrinsic barriers in the maintenance of species boundaries (Abbott *et al.* 2013; Harrison & Larson 2014; Hewitt 1988; Orr & Smith 1998; Turelli *et al.* 2001).

In last few years, next generation sequencing techniques has revolutionized the study of hybridization and speciation processes (for a review, see Seehausen *et al.* (2014)). For instance, recent population genomic studies have provided evidence that adaptive introgression can occur between divergent species and may be more common than previously expected (Abbott *et al.* 2013; Hedrick 2013). The evolutionary histories of the modern human (for a review, see Racimo *et al.* 2015)), the malaria vector mosquito *Anopheles gambiae* (Fontaine *et al.* 2015), *Heliconius* butterflies (Pardo-Diaz *et al.* 2012) and *Mytilus* mussels (Fraisse *et al.* 2016) are particularly well-documented cases illustrating such processes.

Most of these studies are concerned with historical interspecific gene flow which occurred over a long time during periods of range expansion (see Currat *et al.* (2008) for theoretical supports and review of empirical evidences). And yet adaptive introgression may occur on much shorter time scale, as exemplified by introduction of species by human activities which modify species distribution at a global scale and at an unprecedented rate (e.g. in marine ecosystems, see Molnar *et al.* (2008)). Biological introductions provide a window on the early phase of secondary contacts between previously allopatric and non-reproductively isolated species. A diverse set of consequences of hybridization between native and non-native taxa are expected (Allendorf *et al.* 2001) for instance, the extinction of the native species (Rhymer & Simberloff 1996) or the introgression of advantageous alleles from the native into the non-native species facilitating local adaptation of the non-native species to its new colonized environment (Ellstrand & Schierenbeck 2000; Schierenbeck & Ellstrand 2009) or also the opposite situation, i.e. the rapid fixation of non-native alleles in the genome of native species, for example between the non-native Barred Tiger salamanders and the native California one (Fitzpatrick *et al.* 2010).

In this context, we consider two newly reclassified although strongly differentiated species in the genus *Ciona*. These two species were considered as cryptic species of the *Ciona intestinalis* species complex and formerly named *C. intestinalis* type A and *C. intestinalis* type B (Nydam & Harrison 2007; Zhan *et al.* 2010). Following recent taxonomic revision, they are now accepted as two distinct species (WoRMS database) and respectively named *C. robusta* and *C. intestinalis* (Brunetti *et al.* 2015). They display a divergence estimated at ca. 4 Mya (Roux *et al.* 2013). Currently, the two species, and particularly *C. robusta*, display a large distribution over several distinct biogeographic regions because both have been introduced by human-activities (see Supplementary Note in Bouchemousse *et al.* (2016a)). For instance, *C. robusta*, assumed to be native to NW Pacific, has been reported as a non-native species in almost all the oceans. This species lives in sympatry with *C. intestinalis*, native to the NE Atlantic, in only one region, namely in the Western English Channel and South of Brittany. It has been shown that *C. robusta* was introduced in this region probably in the early 2000s (Bishop *et al.* 2015; Nydam & Harrison 2011).

Despite their high divergence (i.e. 14% of transcriptomic divergence (Roux *et al.* 2013) and 12–14% of mitochondrial divergence (Bouchemousse *et al.* 2016a; Nydam & Harrison 2007; Zhan *et al.* 2010)), the two species are not reproductively isolated: first generation (F1) hybrids are easily obtained under laboratory conditions, with however an asymmetry according to the maternal lineage (Bouchemousse *et al.* 2016b; Suzuki *et al.* 2015): F1 hybrids produced in laboratory experiments are obtained in one direction only corresponding to crosses involving oocytes of *C. intestinalis* and sperm of *C. robusta* (ca. 80% of fertilization rate against < 6% in the opposite direction (Bouchemousse *et al.* 2016b)). The question of the extent of hybridization in nature is thus to be addressed. Recent molecular studies carried out in the only sympatric range described so far (i.e. NE Atlantic) suggest contemporary hybridization happens at a small rate: despite a close syntopy and reproductive synchrony, a few putative hybrids (i.e. individuals showing shared alleles on putative species-diagnostic markers) were observed in the wild, with a paucity of F1s (Bouchemousse *et al.* 2016b; Nydam & Harrison 2011). In addition, low levels of introgression were detected and interpreted by the presence of some backcross genotypes in the samples (i.e. between 4 and 6%; Bouchemousse *et al.* 2016b; Nydam & Harrison 2011; Sato *et al.* 2014). If this interpretation is true, this contemporary hybridization could have a profound effect on the expansion of the non-native species but also on the native species (i.e. adaptive introgression processes, see above). However, these studies were based on few nuclear loci assumed species-diagnostic. An alternative explanation is that the low level of admixture measured nowadays in the contact zone is the outcome of historical introgression during past secondary contacts. Roux *et al.* (2013) adjusted a secondary contact model to a population transcriptomic dataset and inferred, under this model of a single contact, that introgression lasted for 15,000 years (95% CI: 4,300 – 56,800) during which ca. 20% of loci presumably crossed the species barriers in both direction. Most probably the two taxa repeatedly came into contacts both in past- and present time. This situation is prone to the misinterpretation of contemporary admixture when few loci are used. Among the samples studied by Bouchemousse *et al.* (2016b), what appear to be a few individuals with hetero-specific alleles could indeed be a consequence of a low genome-wide level of past introgression (e.g. only 1 to 4% of the genome of present-day non-African humans derived from gene flow between Neanderthals and modern humans (Green *et al.* 2010)). In order to ascertain the extent of the contemporary hybridization between the two taxa, we used a population genomic approach based on 310 ancestry-informative SNPs derived from full transcriptomic sequences (Roux *et al.* 2013). By studying such a large number of markers on an extensive sampling, we could also evaluate the discriminating power of the few nuclear markers that have been used so far (e.g. Bouchemousse *et al.* 2016b; Nydam & Harrison 2011; Sato *et al.* 2014). We studied a large number of individuals from eight localities of the sympatric range (i.e. contemporary contact zone) and two to three localities outside contact zones for each species: these allopatric populations were used as a control for the absence of contemporary gene flow between the two species. The SNP panel developed in this study should prove useful in an ascidian species with importance in evolutionary biology, invasion biology, development biology and phylogeny (Procaccini *et al.* 2011; Satoh *et al.* 2014; Zhan *et al.* 2015).

## Materials and Methods

### Sampling

Sampling of *Ciona robusta* and *C. intestinalis* was done within their contemporary sympatric range (i.e. Western English Channel and South of Brittany) in seven localities where the two species are living in syntopy (i.e. living in the same habitat) and one locality where surveys carried out over three years never reported the presence of *C. robusta* (i.e. only *C. intestinalis* is present; no.9 in Table 1 (Bouchemousse *et al.* 2016b)). For comparison, populations from localities outside of the contemporary contact zone (i.e. where a unique species has been recorded so far) were sampled: for *C. robusta*, two localities of the SE Pacific and Mediterranean Sea, and for *C. intestinalis*, two localities in the North Sea (one in shallow water and one at 20-meters depth) and one in the NW Atlantic (Table 1). For each individual, DNA extraction was performed with Nucleospin^®^ 96 Tissue Kit according to the manufacturer’s protocol (Macherey-Nagel, Germany). A minimum of 24 individuals per population was selected based on the DNA quality following extraction. Altogether a total of 449 individuals, 213 for *C. robusta* and 236 for *C. intestinalis* were further analyzed. A preliminary assignment to either *C. robusta* or *C. intestinalis* was based both on morphological features (Brunetti *et al.* 2015; Sato *et al.* 2012). In addition to specimens sampled in natural populations, two F1-hybrids produced from experimental crosses (Bouchemousse *et al.* 2016b) were included as control for F1-hybrid genotype.

**Table 1.**
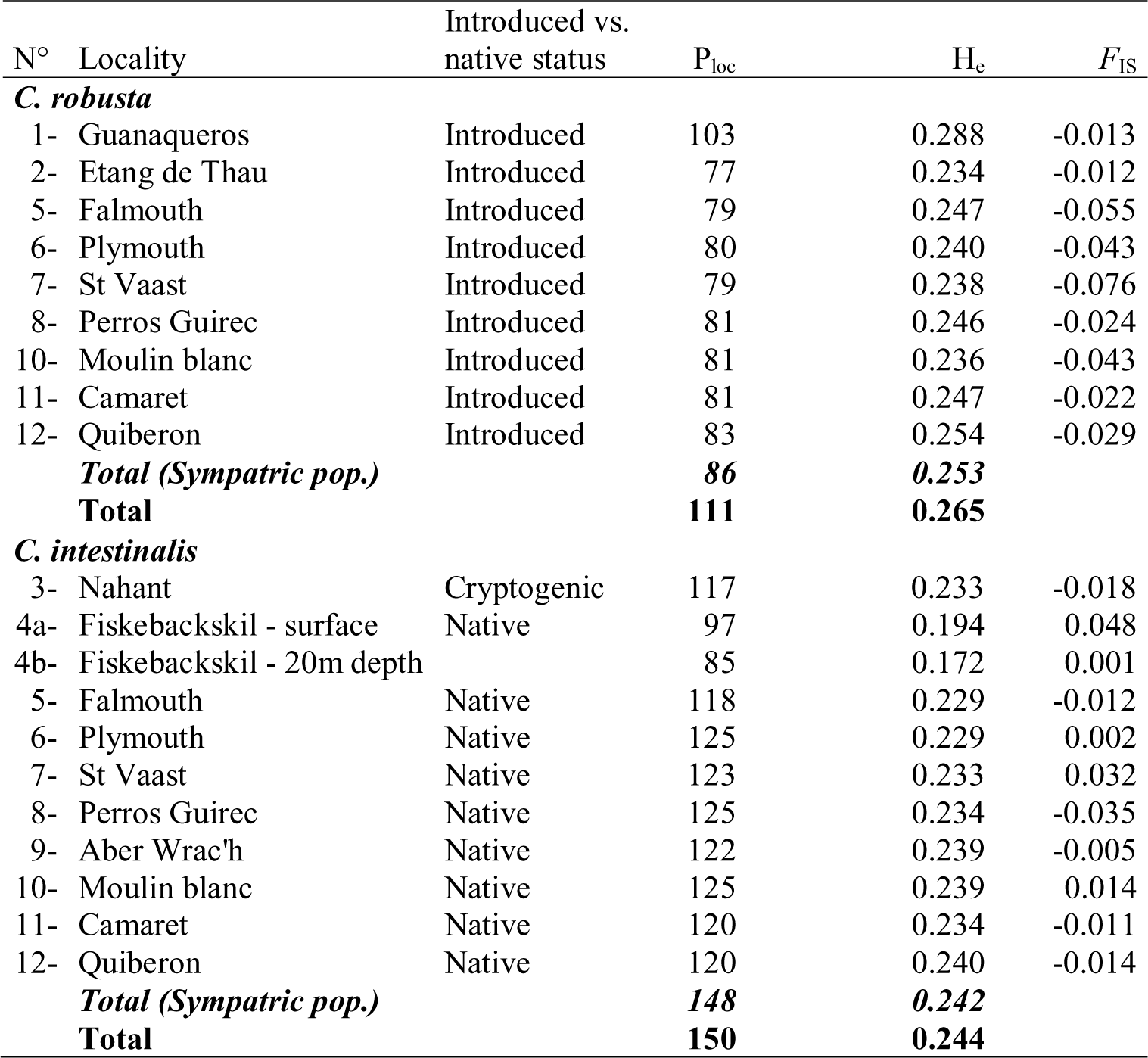
**Study localities, hybrid index (*h*) and number of hybrids *sensu lato*** (i.e. F1, F2 hybrids and backcrosses with parental species according to NEWHYBRID software) **in each population of *Ciona robusta* and *C. intestinalis***. *Regional status* and *locality status* indicate if the two species have been reported to co-exist at a regional scale (allopatric vs. sympatric) or at the locality level (syntopic vs. non-syntopic). In this table, *h* is defined as the proportion of alleles from one species in the genetic background of the other species (i.e. proportion of *C. intestinalis* alleles over all loci in *C. robusta* individuals and proportion of *C. robusta* alleles over all loci in *C. intestinalis* individuals). *h* values were averaged across individuals for each sampled localities. Analyses were done with 105 SNPs selected for inter-specific gene flow analyses (*F*_ST_ > 0.9; see Material and Methods).

### Loci selection and genotyping

An Illumina BeadXpress^®^ with Veracode^TM^ technology (GoldenGate^®^ Genotyping Assay) was used to genotype 384 single nucleotide polymorphisms (SNPs) selected from a SNP dataset detected in the full transcriptomes of 10 individuals of *C. robusta* and 10 individuals of *C. intestinalis* (details in Roux *et al.* (2013)). The loci were first chosen to maximize their genotyping success: Because we used transcriptome data, we identified exon borders by aligning our data with *C. robusta* genome (vKH.71 from Ensembl; note that the genome name is misleading as it is labelled “*C. intestinalis*” although it is from *C. robusta* following the recent taxonomic revision (Brunetti *et al.* 2015)). Polymorphic sites closer than 20bp from exon border were automatically excluded. Polymorphic positions were selected within exons to produce an individual sequence for each given SNP compatible with the Assay Design Tool (ADT) software available on Illumina webpage. Sites with a minor allele frequency lower than 0.1 were excluded. ADT software was used to choose primers for each SNP and estimate probability of amplification of each marker before amplification. Only markers with a probability of amplification greater than 40% were retained. We selected this low minimum value because of the high divergence between *C. robusta* and *C. intestinalis* at the full transcriptome level (i.e. 14% according to Roux *et al.* (2013) and thus the poor number of genomic regions likely to be conserved between the two species. The average probability obtained for our final SNP panel of 384 markers was however reaching 74%. Based on the results by Roux *et al.* (2013), loci could be sorted according to four categories of polymorphism (Table S1): 1) SNPs differentially fixed between the two species (sf), 2) SNPs polymorphic in *C. robusta* (sxA) but not in *C. intestinalis*, 3) SNPs polymorphic in *C. intestinalis* (sxB) but not in *C. robusta* and 4) SNPs displaying polymorphism in the two species (ss). The full SNP panel was intentionally not random, for instance including a substantial number of SNPs differentially fixed between the two species and shared polymorphisms when compared to the genome-wide expectation. Selecting loci showing a high genetic differentiation between reference populations, such as allopatric populations located far away of a sympatric zone or from either side of a hybrid zone is common practice for discriminating recent admixed individuals (Bierne *et al.* 2011; Larson *et al.* 2014). However, these loci display restricted introgression and might not randomly associate during reproduction (Harrison & Larson 2016). Thus, we expanded the SNP panel with polymorphic loci showing less divergence and shared between the two species. In addition, the 384 SNPs were selected to be spread over most of the chromosomes of the published genome of *C. robusta* (Dehal *et al.* 2002) and 25 of them were localized in introgression hotspots identified by Roux *et al.* (2013)). At the end, we enriched the panel with sf (101) and ss (47) SNPs and equalized the number of sxA (109) and sxB (127) SNPs as *C. intestinalis* is more polymorphic than *C. robusta*. A subset of 70 SNPs that strictly reflect the genome wide site frequency spectrum (i.e. randomly selected) was included in the SNP panel. Genotyping was performed using Genome Studio software (Illumina Inc.). Out of the 384 SNPs, 324 SNPs amplified successfully and 310 SNPs were retained for further statistical analyses. Despite the expected low amplification score (probability of amplification *in silico* of 74%), these 310 SNPs displayed a high rate of genotyping success: we obtained a minimum of 97% of the individuals without missing data, and an unambiguous genotype assignment. This SNP panel included 58 SNPs randomly selected over the initial subset of 70 SNPs (the remaining 12 SNPs have not been successfully genotyped).

In order to investigate the properties of putative species-diagnostic markers used in previous studies (Bouchemousse *et al.* 2016b, Caputi *et al.* 2007, Nydam & Harrison 2010, 2011, Sato *et al.* 2014), we also genotyped all the individuals on three nuclear loci, namely Hox5 (Caputi *et al.* 2007 vAChTP and CesA (Nydam & Harrison 2010) by PCR and PCR-RFLP (details in Caputi *et al.* (2007)) and Nydam & Harrison (2010)) and a putative maternal species-diagnostic mitochondrial locus (mtCOI; details in Nydam & Harrison (2007)). Details regarding the physical mapping of the nuclear loci are provided in Figure S1 and Table S2.

### Intra-specific analysis

In order to compare sympatric and allopatric populations, genetic studies were carried out for both *C. robusta* and *C. intestinalis*. Only loci that were polymorphic in the targeted species were used. For the few SNPs chosen in the same contig, only one SNP was selected, the one showing the maximum value of the minor allele frequency. Totals of 111 and 150 polymorphic loci were retained for *C. robusta* and *C. intestinalis*, respectively.

#### Genetic diversity

At the intra-specific level, for each population, the number of polymorphic loci and the expected heterozygosity (*H*_*e*_) were estimated using GENETIX v.4.05 software (Belkhir *et al.* 2004). Fixation index (*F*_IS_) was estimated and departures from Hardy-Weinberg equilibrium were tested in each population using GENEPOP v4 with default parameters for tests. *P*-values resulting of multiple tests were adjusted using the R package v. 3.1.3 (R Development Core Team 2014) QVALUE (Storey 2002).

#### Genetic structure

Genetic structure between populations was analyzed by estimating the fixation index *F*_ST_ (Wright 1951) using GENEPOP. Exact G test for population differentiation were carried out using 10,000 random permutations. To visualize the genetic structure between populations, a Discriminant Analysis of Principal Components (DAPC; (Jombart *et al.* 2010)) was computed for each species separately using the R package ADEGENET v.1.4 (Jombart & Ahmed 2011).

### Genome and population admixture analysis between the two species

To identify putative F1- or recently introgressed individuals (product of several generations of backcrosses within the sympatric range), a Bayesian clustering method implemented in NEWHYBRID software v.1.1 was used (Anderson & Thompson 2002) using the global dataset of 310 loci, the dataset of 58 random SNPs, as well as a dataset of the 105 most differentiated loci. Briefly, this method computes posterior probability for assigning a given individual to different hybrid categories (i.e. F1, F2-hybrids and backcrosses with parental *C. robusta* or *C. intestinalis* individuals) or parental species, using Markov chain Monte Carlo algorithms (MCMC). Here, we considered allopatric populations as representative of the parental species. We ran five independent analyses (each using a different random starting value) with 500,000 MCMC after a period of 500,000 burn-in cycles using the Jeffreys-like prior.

To examine inter-specific gene flow at the genome level, we used the R package INTROGRESS (Gompert & Buerkle 2010). As the ancestral allelic state has to be defined (based on the minor allele frequency in each species), we selected 105 loci that were the most differentiated loci according to *F*_ST_ values (*F*_ST_ > 0.9) computed between the two species using allopatric populations (Figure S2). Such a selection is common practice in hybridization studies (e.g. between *Mytilus* species (Saarman & Pogson 2015) or *Gryllus* crickets (Larson *et al.* 2013, 2014)). For each individual, the maximum likelihood value of hybrid index was estimated. The hybrid index (*h*) is defined as the proportion of *C. intestinalis* alleles over all loci (*h* = 0 for individuals with *C. robusta* alleles only and *h* = 1 for individuals with *C. intestinalis* alleles only). To visualize and compare the genomic architecture of interspecific admixture at the individual level, the *mk.image* function implemented in INTROGRESS was used.

NEWHYBRIDS and INTROGRESS analyses were also done using a small dataset made of the three nuclear markers used in previous studies as putatively species-diagnostic (i.e. Hox5, vAChTP and CesA, see “*Loci selection and genotyping*” section above).

The inter-specific admixture rate was investigated on the total dataset (i.e. 310 loci) using a Bayesian clustering method implemented in STRUCTURE v.2.3 (Pritchard *et al.* 2000) and a Principal Component Analysis (PCA) using the R package ADEGENET v.1.4 (Jombart & Ahmed 2011). The method implemented in STRUCTURE method used MCMC to generate posterior probabilities of assignment of each individual genotype to a number of clusters (K). Ten replicates of 500,000 MCMC after a period of 500,000 burn-in cycles were ran for K values ranging from 1 to 4; K = 2 is corresponding to the two species clusters. Results were summarized across all replicate runs using CLUMPP v.1.1.2 (Jakobsson & Rosenberg 2007) and visualized with DISTRUCT v1.1 (Rosenberg 2004). To check for the absence of biases due to marker selection, we also ran STRUCTURE for K = 2 using additional dataset: 1) a subset of 58 loci selected to represent a random sampling of the genome, 2) a subset of 105 loci selected for the INTROGRESS analysis, 3) 245 SNPs corresponding to all the SNPs except those differentially fixed between the two species, 4) only those SNPs that were polymorphic in the two species (42 SNPs).

To better evaluate the evolutionary history between *C. robusta* and *C. intestinalis* populations, we used a population graph approach implemented in the TREEMIX program (Pickrell & Pritchard 2012), which infers patterns of splitting and migration between populations. By using the matrix of allele frequency covariance between pair of populations, this method generates maximum likelihood population trees under the hypothesis of an absence of migration or the alternative hypothesis of migration event(s) (that are sequentially added). By comparison with other methods commonly used to make demographic inferences or to picture population relationships over time (e.g. Beaumont *et al.* 2002; Gutenkunst *et al.* 2009; Hey & Nielsen 2001), TREEMIX has the advantage to be applicable to a large number of populations by using a tree-construction based approach and, at once, testing for gene flow between populations (Pickrell & Pritchard 2012). To avoid noises due to small sample sizes and intra-specific migration (i.e. infra-specific admixture), we pooled populations according to the region of sampling (i.e. no.4a and 4b for *C. intestinalis*; no.5 and 6 and no. 7 to 12 for each species). Using the total dataset (i.e. 310 loci), we search for the best tree to fit the data testing for a range of migration events from 0 to 8 (reaching an asymptotic value, Figure S3). Based on these inferences, we used a block-jackknife procedure with blocks of 10 SNPs to determine which migration events significantly improved the model fit. A complementary analysis based on *f*_3_-statistic test, developed by Reich *et al.* (2009), was done to test the null hypothesis that the evolutionary history of *Ciona* populations was consistent with the absence of migration events between populations. The *f*_3_-statistics evaluates the deviation of the null hypothesis using the same block-jackknife procedure for all combinations of three populations (one used as the target and two tested as putative ancestral populations).

## Results

### Diversity of the SNP panel

Overall, 451 individuals (including the two F1-hybrids from experimental crosses) were genotyped successfully at 310 SNPs defined from a transcriptome dataset of *Ciona robusta* and *C. intestinalis* (Roux *et al.* 2013). Following this genotyping, the distribution of SNPs across the categories, defined from a small sample of 10 specimens for each species, was modified, as shown in Table S1. The most substantial change was a decrease of the sf and sxA categories (i.e. a decrease of 31% and 22%, respectively) and a concomitant increase of the sxB and ss categories (17% and 22%, respectively). We considered these new categories in the analyses below.

### Population genetic structure and little heterozygosity variations in the two study species

The analyses aiming at comparing allopatric and sympatric populations of each species were carried out separately for *C. robusta* and *C. intestinalis,* using the set of loci polymorphic in each species, i.e. 111 and 150 SNPs respectively. Results are summarized in Table 2 and Table 3.

**Table 2.**
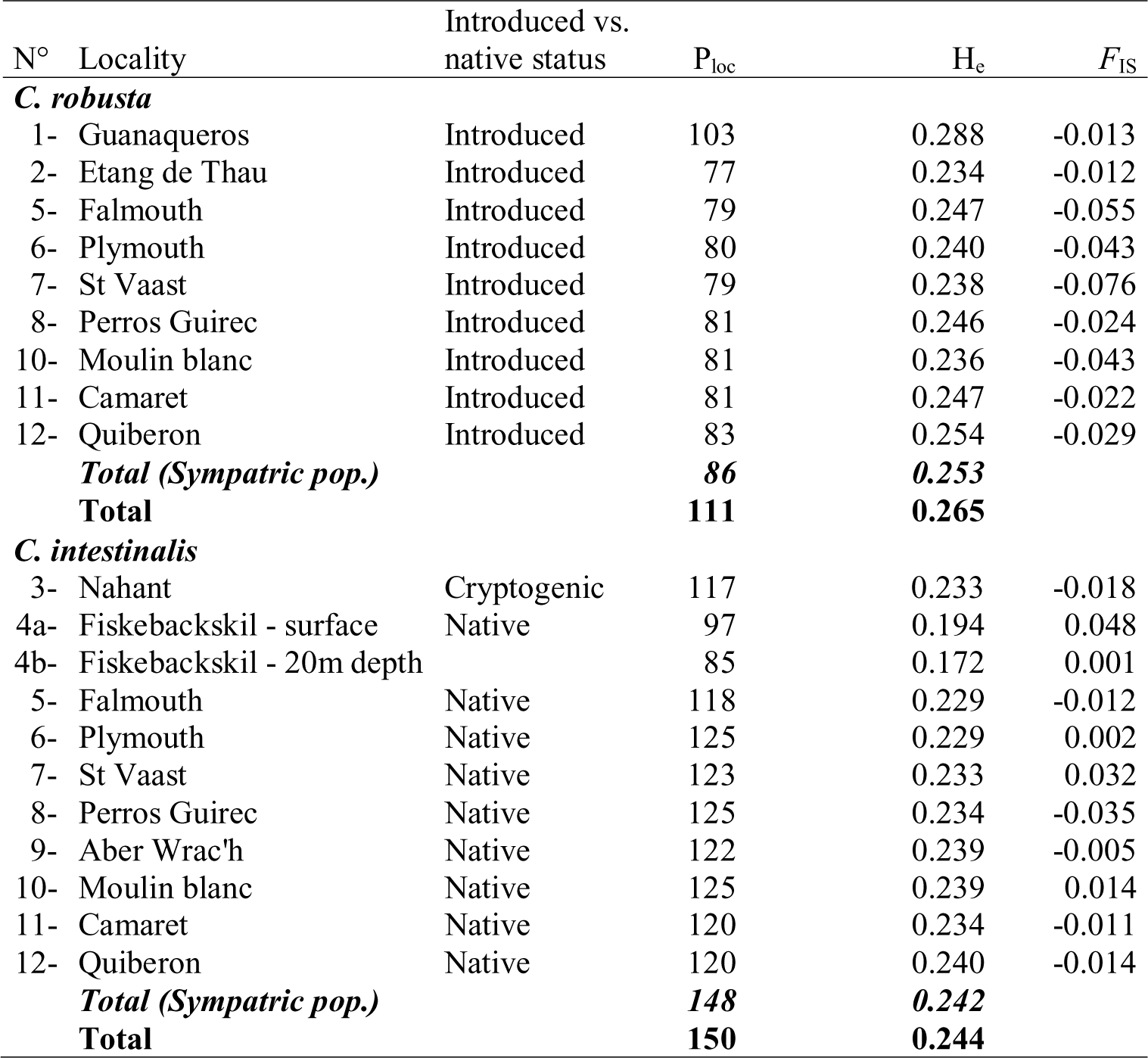
**Genetic diversity indices and fixation index** of each study populations for *Ciona robusta* and *C. intestinalis*. P_loc_: number of polymorphic loci and H_e_: expected heterozygosity over 111 and 150 polymorphic loci retained for intra-specific analyses in *C. robusta* and *C. intestinalis,* respectively (see *Materials and Methods*); *F*_IS_: fixation index calculated (no deviation from Hardy-Weinberg equilibrium; exact test, *P* < 0.05).

**Table 3.** Genetic structure among populations for *Ciona robusta* and *C. intestinalis*.

| | $F_{ST}$ | $P$-value |
| --- | --- | --- |
| ***C. robusta*** |  |  |
| All sampled populations (9 populations) | 0.054 | $P < 0.001$ |
| All populations without Guanaqueros (all except no.1) | 0.023 | $P < 0.001$ |
| Sympatric populations (all except no.1 and 2) | 0.021 | $P < 0.001$ |
| ***C. intestinalis*** |  |  |
| All sampled populations (11 populations) | 0.045 | $P < 0.001$ |
| All populations without Fiskebackskil (all except no.4a and 4b) | 0.021 | $P < 0.001$ |
| Sympatric populations (all except no.3, 4a and 4b) | 0.014 | $P = 0.020$ |

#### *Diversity and genetic structure in populations of* C. robusta

Values of *He* were similar across populations of *C. robusta*, ranging from 0.234 (no.2) to 0.288 (no.1). No departure from Hardy-Weinberg equilibrium (HWE) was found in any of the study populations. Exact test of differentiation revealed significant differences in allele frequencies among all populations sampled and among populations of the sympatric range (Table 3). The highest genetic differentiation was observed between the SE Pacific population (no.1) and all of the other populations (pairwise comparisons provided in Table S3a) as well-illustrated by the DAPC (Fig.1a) along the first discriminant axis. The second discriminant axis pointed out the differentiation between populations of UK (i.e. no.5 and 6) and Mediterranean Sea (no.2) which is confirmed by significant pairwise estimates of *F*_ST_ (Table S3a). Populations of Brittany were relatively poorly differentiated between them (non-significant *F*_ST_ values in most of pairwise comparisons, Table S3a). Altogether, SE Pacific and to a lesser extent UK and Mediterranean Sea populations were the most different genetically.

**Figure 1.**
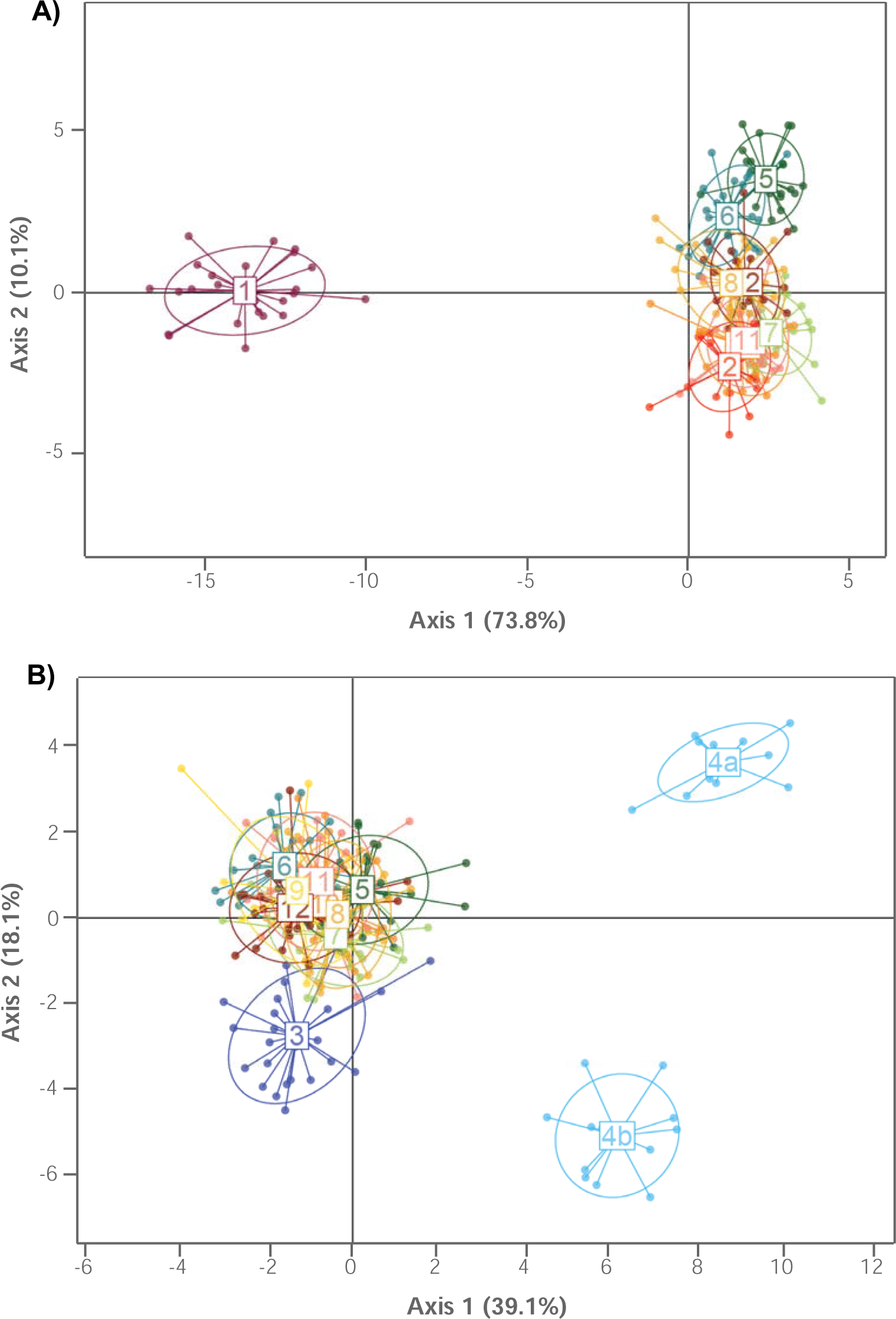
Discriminant Analysis of Principal Components (DAPC) among populations of *C. robusta* (A) and *C. intestinalis* (B). Only the two first axis showing the two higher discriminant eigenvalues are presented here.

#### *Diversity and genetic structure in populations of* C. intestinalis

Values of *He* were similar among the study populations, ranging from 0.240 (no.12) to 0.229 (no.5 and 6), except for the populations from the North Sea which exhibited lower values of *He* (i.e. 0.194 and 0.172 for no.4a and 4b respectively). As for *C. robusta*, no departure from HWE was observed in any study populations. Exact test of differentiation between *C. intestinalis* populations indicated significant differences among all populations but was non-significant between populations of the sympatric range (Table 3). The overall significant genetic structure was mainly due to a strong and significant genetic differentiation of the populations sampled in the two allopatric regions (no.3, 4a and 4b) and of one population sampled in the sympatric range (no.6) with almost all other populations (pairwise comparisons are provided in Table S3b). These patterns are pictured by the DAPC (Fig.1b).

### Low hybrid index disregarding the regional category and population status

A total of 105 loci, showing a *F*_ST_ strictly superior to 0.9, were used with the R package INTROGRESS to examine the patterns of shared polymorphism between the two species in the contact zone and in allopatric populations. At the species level (i.e. across all individuals for each species), values of *h* were very low, with an average value across individuals of 0.0029 for *C. robusta* and 0.0055 for *C. intestinalis*. Table 1 is providing the average values of the hybrid index (*h*) for each population of *C. robusta* and *C. intestinalis*. A noticeable result was the presence of one individual in this latter population with an *h* value of 0.5. When removing this individual from the *h* estimation, the value in Camaret dropped to 0.006, a value close to the average values for *C. intestinalis* populations. This individual was assigned with a probability of 1 to a ‘F1 hybrid’ with NEWHYBRIDS (Table 1) with the 105 SNPs dataset; a result confirmed with the full (310 SNPs) and the random (58 SNPs) dataset. The two F1-hybrids obtained experimentally were also assigned with a probability of 1 to the ‘F1-hybrid’ category which ascertains the robustness of NEWHYBRIDS for detecting individuals derived from recent crosses. It is noteworthy that all the other study individuals were assigned to their respective parental ‘species’ categories (Table 1). We also examined the relationship between *h*-value and the heterozygosity rate across the 105 loci used with INTROGRESS, by using a triangle plot displayed in Figure 2: all except one individual displayed extreme *h*-values (closed to 0 or 1) and an extremely low proportion of heterozygote loci for *C. robusta* and *C. intestinalis* alleles. The only exception is the individual sampled from Camaret (no.11) that was assigned by NEWHYBRIDS as a F1-hybrid: this individual showed both a high *h*-value and a high heterozygosity rate (i.e. 99%); these values were similar to the values observed for the two F1-hybrids experimentally produced in the laboratory (Fig.2, i.e. 96% and 99%). STRUCTURE analyses with K = 2 assigned equally the putative wild F1-individual and the two experimental F1-hybrids to the two species clusters (Fig.3b); and the results of the PCA showed a clear distribution of the overall genetic variance between the two study species with the natural and experimental F1-hybrids at an intermediate position (Fig.3a). This finding is also observed on STRUCTURE analyses done with the additional subset of loci (Figure S4; see *Material and Methods* section) suggesting an absence of biases due to marker selection. Note that increasing K values (K = 3 and K = 4, Fig.3b) in the STRUCTURE analysis confirmed intraspecific variance observed with DAPC (Fig.1), notably the genetic differentiation of the SE Pacific population (no.1) with all of the other populations of *C. robusta* and of the two sub-populations of Fiskebackskil with all of the other populations of *C. intestinalis*.

**Figure 2.**
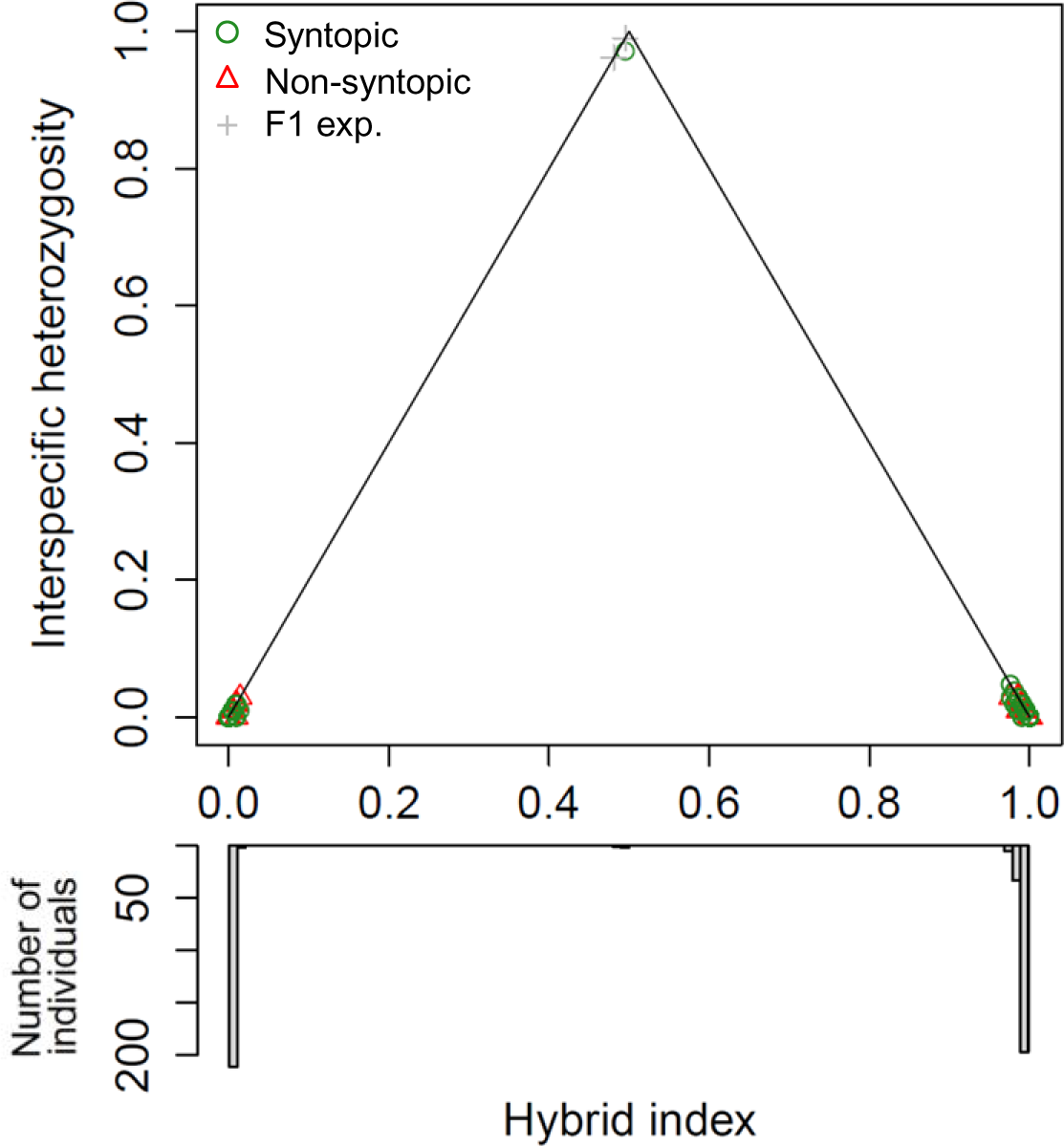
Triangle plot showing the relationship between heterozygosity rate across loci and hybrid index for each individual. At the top of the triangle, one green circle is picturing one individual from the locality no. 11 and the two gray crosses are F1-hybrids from experimental crosses.

**Figure 3.**
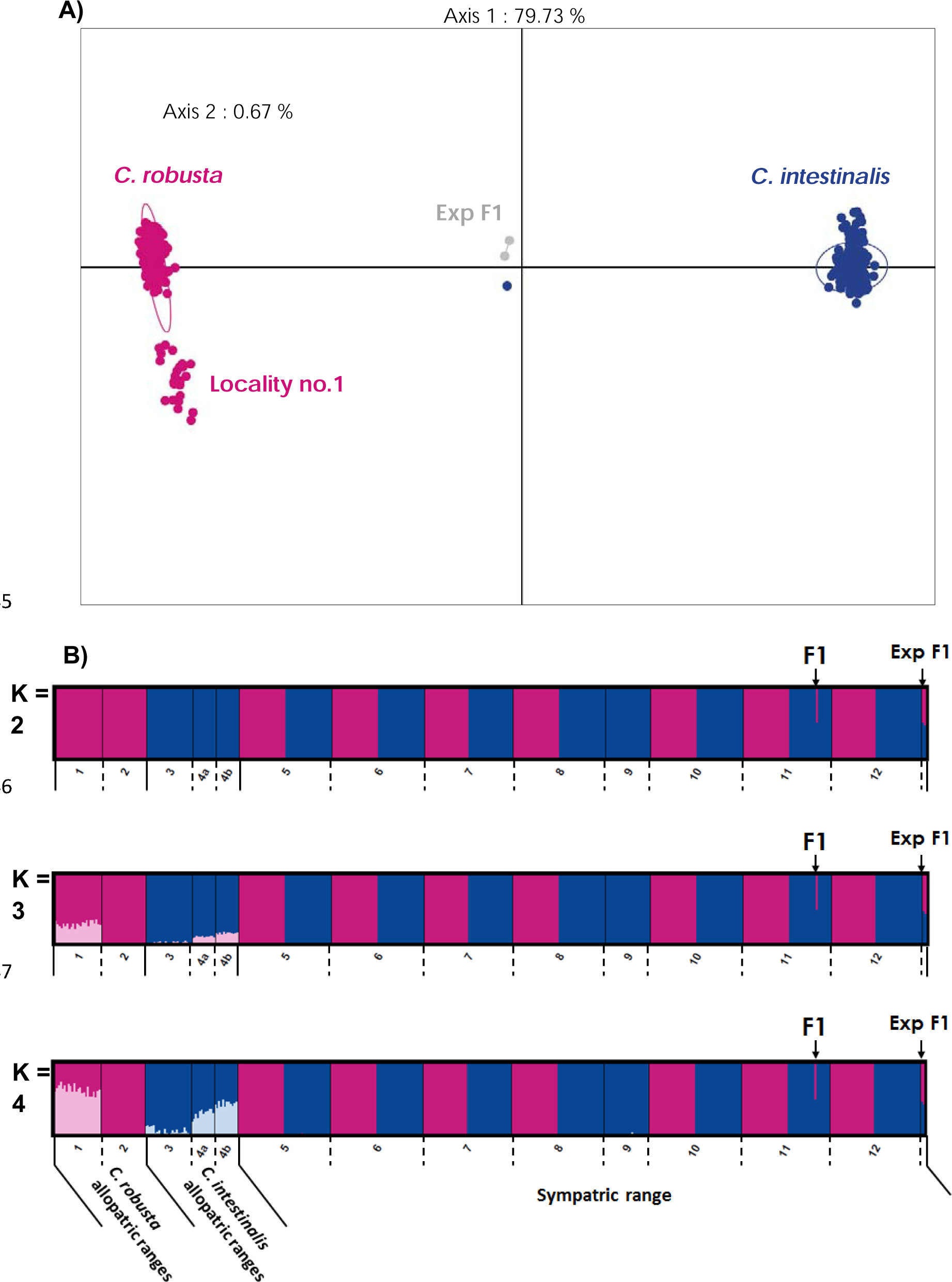
A) Principal Components Analysis (PCA) and B) Individual Bayesian assignment proportion for clusters from K =2 to K = 4. using the total dataset (i.e. 310 SNPs, 449 individuals from natural population and the two F1 hybrids from experimental crosses).

Using the three nuclear markers used as species-diagnostic markers in previous studies (i.e. Hox 5, vAChTP and CesA; see Material & Methods), an interesting pattern was observed: the single F1 individual otherwise identified with the complete set of SNPs displayed a heterozygote genotype at two loci (i.e. vAChTP and Hox5) but a homozygote genotype (two *C. robusta* alleles) at CesA locus; a pattern clearly inconsistent with a F1-hybrid genotype if loci are assumed fully diagnostic. In addition, NEWHYBRIDS assigned with a high probability (*P* = 0.95) this F1-hybrid individual to the category of individuals backcrossed with *C. robusta*. This result illustrates the inherent difficulty to accurately account for the sampling variance when alleles are fixed in the samples or nearly so. Among the other individuals, 416 (92.8%) were assigned to their respective parental ‘species’ categories with a probability above to 0.95, while 32 individuals (7.2%) obtained ambiguous results with a posterior probability of being parental genotypes ranging from 0.63 to 0.95. At the mtCOI locus (i.e. a putative maternal species-diagnostic mitochondrial locus), there is a strict association between the preliminary morphological assignment and the mitochondiral type (Fig.4b). As expected based on experimental studies (i.e. asymetry in reproductive success according to the maternal type, see Introduction section), the F1 hybrid showed a *C. intestinalis*-mitochondrial type.

**Figure 4.**
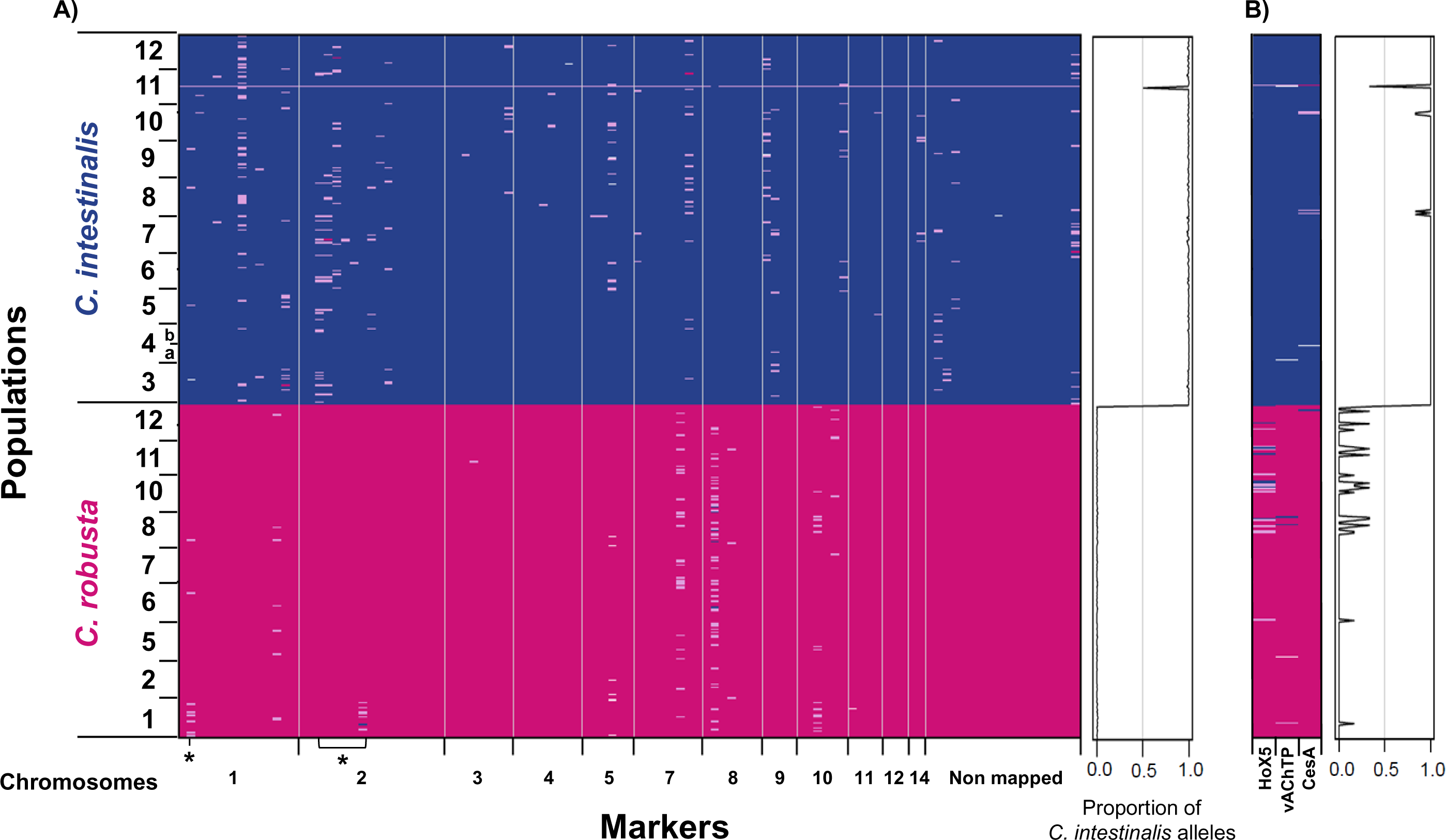
A) Genomic architecture using 105 highly differentiated loci (*F*_st_ > 0.9) selected for inter-specific analyses. Markers (x-axis) are ord following physical position on chromosomes. Individuals (y-axis) are ordered per population. Dark pink cases indicate homozygote genotype on *C. rob* alleles; dark blue, homozygote genotype on *C. intestinalis* alleles; light purple, heterozygotes for *C. robusta* and *C. intestinalis* alleles; and white cases, mis values. Asterisks indicate loci located in introgression hotspots defined by Roux *et al.* (2013). **B) Pattern of admixture for 3 nuclear loci** (Hox5, vACh CesA) **and one mitochondrial locus** (mtCOI) **analyzed by PCR and PCR-RFLP**, already used in previous studies (Nydam & Harrison 2011; Sato *et al.* 2014; Bouchemousse *et al.* 2016).

### Heterogeneous polymorphism rates at the genome level

The subset of 105 SNPs used for examining the patterns of shared polymorphism between the two species (i.e. showing *F*_ST_ values higher than 0.9) showed the following patterns over the whole dataset: 65 were differentially fixed (i.e. sf loci), 39 with private polymorphisms (i.e. sxA or sxB; 8 polymorphic in *C. robusta* and 31 in *C. intestinalis*) and only one locus showing shared polymorphism (i.e. ss locus) between populations of the two species (snp18 on the chromosome 1). The 40 loci showing shared and private polymorphisms were distributed randomly along the genome of the two species (Fig.4a), as previously observed in Roux *et al.* (2013). Among the 105 SNPs, some were found in two introgression hotspots defined by Roux *et al.* (2013). Note that we did not have SNPs localized in the other two introgression hotspots. Interestingly, we found shared or polymorphic SNP in these introgression hotspots: 1) one SNP in the introgression hotspot of chromosome 1 showed shared polymorphism in the two species, and 2) six loci showed private polymorphism in one or the other of the two species in the introgression hotspot on the chromosome 2.

Polymorphism patterns were also informative regarding the status of the study populations (i.e. allopatric and sympatric). Details of allele frequencies at each of the 40 polymorphic loci in the allopatric and sympatric ranges of the two species are provided in Table S4. When comparing of populations for *C. robusta* and *C. intestinalis*, the rates of shared and private polymorphism appeared to be remarkably stable across populations: for the two species, individuals of each population carried a small number of heterozygous sites, but not always at the same genome location (Fig.4a). A noteworthy exception was the allopatric population of *C. robusta* from Chile (no.1) which shared for some loci more polymorphism with *C. intestinalis* populations than with other populations of *C. robusta* (Fig.4a): heterozygous sites were more important, for example, at the snp18 (chromosome 1), snp290 (chrom. 2) and snp237 (chrom. 10), the two first being in introgression hotspots defined by Roux *et al.* (2013). This finding was already visible in the results of the PCA (Fig.3a) as the population of Chile was slightly shifted towards the *C. intestinalis* points

The random distribution of few shared and private polymorphisms was also observed when using the three putative species-diagnostic markers (Fig.4b): minor allele frequency observed for Hox5, vAChTP and CesA was 0.2, 0.2 and 1.3% for *C. intestinalis*, and 6.4, 1.2 and 0.5% for *C. robusta*, respectively. None of them were localized in introgression hotspots. As for the 40 polymorphic SNPs discussed above, the rate of polymorphism was quite stable across populations with a small number of heterozygous sites.

### Admixture events between the two species revealed by a population tree approach

The population tree inferred from TREEMIX without migration explained 88.5% of the variance in the population covariance matrix. Note that in the population tree without migration events, the population of Chile (no.1) showed a position shifted towards *C. intestinalis* populations (Figure S5). The variance explained was increased when migration events were added (Figure S3). The best fit to the data was obtained with two migration events, which significantly improved the model (*P* < 0.001, Fig.5). This population tree, explaining 98.5% of the variance (Figure S3), indicated significant gene flow in *C. robusta* population of Chile (no.1) and in the *C. intestinalis* populations group of Brittany (no.7 to 12). These two migration events were also supported by the *f*_3_ statistics analysis (Table S5) with significant negative values for almost all combinations of three populations involving as targets the *C. robusta* population of Chile (no.1) and the *C. intestinalis* populations group of Brittany (no.7 to 12). *f*_3_ statistics also showed significant negative values for combinations of three populations involving as targets *C. robusta* populations groups of UK (no.5 and 6) and Brittany (no.7 to 12) (Table S5). These negative *f*_3_ statistics are consistent with the hypothesis that the tested populations were the results of admixture with ancestors in the two tested population sources (Reich *et al.* 2009).

**Figure 5.**
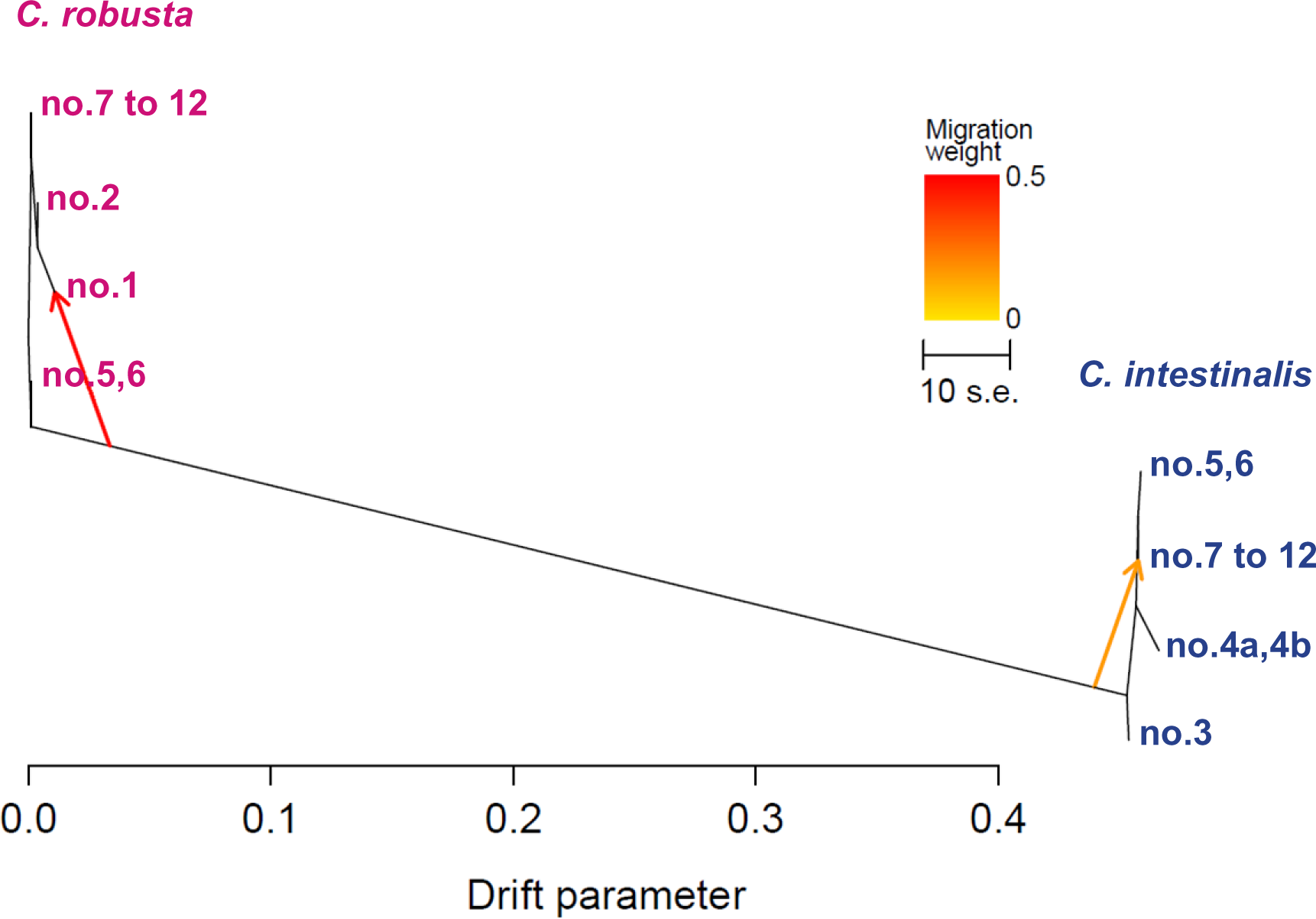
Population tree inferred by TREEMIX indicating two migration events between *Ciona robusta* and/or *C. intestinalis* populations using the total dataset. (i.e. 310 SNPs). Terminal nodes are labelled by locality number (Table 1). Note that we pooled populations according to regions of sampling (i.e. no.4a and 4b for *C. intestinalis*; no.5 and 6 and no. 7 to 12 for each species) to avoid noises by intra-specific admixture events. Admixture arrows are colored according to the migration weight. The two admixture events significantly improved the model as compared to a situation without migration (*P* < 0.001).

## Discussion

In this study we used 310 ancestry-informative SNPs to clarify relative contribution of contemporary hybridization *versus* past introgression in the level of shared polymorphism observed between *Ciona robusta* and *Ciona intestinalis,* and to analyze the introgression patterns within allopatric and sympatric ranges of the two species. These two points are discussed in turn below.

### Absence of contemporary interspecific gene flow in the sympatric range

In previous studies that analysed interspecific gene flow in the sympatric range, admixture have been observed between the two species although at low rates: 4.2% (including one putative F1-hybrid) over 730 individuals sampled in Nydam & Harrison (2011), 6.3% over 288 individual sampled in one locality by Sato *et al.* (2014), 4.3% (including one putative F1 hybrid) over ca. 3,000 individuals by Bouchemousse *et al.* (2016b). For examining the extent of hybridization between the two species, these authors used few nuclear markers (between 3 and 6 loci according to the study) which were supposed to be species-diagnostic. Consequently, discriminating the footprint left by historical introgression *versus* contemporary hybridization was particularly difficult as acknowledged in these studies (Bouchemousse *et al.* 2016b; Nydam & Harrison 2011). Using a large number of informative loci with a large sample, we found that with the exception of a single individual all the other occurrence of shared polymorphism are the likely consequence of a low level of past introgression, i.e. ancient gene flow between the two species, rather than contemporary hybridization. At a given locus some individuals can be found heterozygotes at quasi diagnostic loci, but averaging at many such loci shows every individual have the same hybrid index value. The only contemporary hybrid was a F1, as supported by both NEWHYBRIDS (whatever the dataset used) and INTROGRESS analyses (Table 1, Fig.2 and 4a). The mtDNA type of this individual is typical of *C. intestinalis* which corresponds in many studies to crosses easily produced in laboratory experiments (Bouchemousse *et al.* 2016b; Suzuki *et al.* 2005). The presence of one F1 hybrid only in our study confirms the hypothesis by Sato *et al.* (2014) and Bouchemousse *et al.* (2016b) of the existence of pre-zygotic isolation mechanisms preventing contemporary hybridization in the wild. However, our new interpretation that recent backcrosses of a few generations are completely lacking from the sympatric range, suggest that strong post-zygotic selection is also occurring, which is the least one can expect for two highly divergent species (i.e. 14% of divergence based on transcriptomic data (Roux *et al.* 2013)). Dobzhansky-Muller incompatibilities expressed by recessive mutations in subsequent generations of hybridization (e.g. Bierne *et al.* (2006); Fishman & Willis (2001) and see for a review Maheshwari & Barbash (2011)) are likely the cause of this isolation. Altogether these results confirm that contemporary gene flow is almost inexistent between the two species.

### The footprint of past introgression between the two species

Hybrid index and interspecific heterozygosity values were similar whatever the region (sympatric or allopatric) and locality status (syntopic vs. non-syntopic, Table 1, Fig. 2). This finding validates that shared polymorphism were observed in all populations including in localities of allopatric regions (Fig.4a). Footprints of gene flow were also observed and were significant for some of them according to TREEMIX and *f*_3_-statistics analyses.

For a given locus, shared polymorphism or high derived allele frequency between two species may result from incomplete lineage sorting of ancestral polymorphism, contemporary or past secondary introgression, or homoplasy. In the case of the two *Ciona* species studied here, the contemporary introgression hypothesis can be reasonably excluded as discussed above. Concerning incomplete lineage sorting of ancestral polymorphism, it would have meant that the polymorphism observed nowadays would have been maintained randomly across loci after the allopatric divergence estimated to have occurred during the Pliocene (between 2.59 and 5.33 My (Roux *et al.* 2013)). Considering the long time elapsed since the divergence, the probability of occurrence of the two ancestral alleles in both daughter species is likely to be extremely low under a neutral model (Pamilo & Nei 1988). High effective population sizes moderates the effect of genetic drift and so the probability of fixation of alleles over the time (Maddison 1997; Pamilo & Nei 1988). *Ciona* species and their common ancestor were characterized by high effective population sizes, estimated in Roux *et al.* (2013), as between 115,000 and 395,000 for *C. robusta*, 748,000–1,022,000 for *C. intestinalis* and 1,606,000–2,220,000 for the common ancestor. However, the analysis of Roux *et al.* (2013) showed that the strong excess of shared polymorphism between the two species cannot be obtained without secondary introgression. The secondary contact has been estimated to have occurred 15,500 years ago (95% CI: 4,300–56,800), during which ca. 20% of loci crossed the species barrier in both directions. Besides similarities in admixture levels across localities, the hypothesis of an ancient admixture event is also well-supported 1) by significant admixture events between populations of *C. robusta* and *C. intestinalis* according to TREEMIX and *f*_3_-statistics analyses and 2) the presence of admixed loci in introgression hotspots (i.e. loci pointed by an asterisk in Fig.4a).

Our finding is also interesting to consider in light of previous studies based on a few markers used as species-diagnostic markers of the two study species (e.g. (Bouchemousse *et al.* 2016b; Nydam & Harrison 2011; Sato *et al.* 2014)). Analyzing a small number of such loci can easily result in the erroneous interpretation that some individuals are more admixed than other and cast doubts about the ability of these markers to reliably distinguish the two species. This was shown in our study by comparing the results obtained with 310 SNPs vs. three markers supposed to be species-diagnostic. In particular, the CesA locus showed a homozygote genotype with two *C. robusta* alleles in the single F1 individual otherwise identified with the complete set of SNPs. With the subset of three markers, this F1-hybrid individual was consequently mistaken as a backcrossed individual and not assigned to the F1-hybrid category, with NEWHYBRIDS.

These results highlight the risks of using putative species-diagnostic markers without preliminary knowledge about the likelihood of past introgression between two study taxa. The species complex of *Mytilus* species is another well-known case study: *Glu* and *mac-1* loci were mistakenly considered as diagnostic makers for *M. galloprovincialis* and *M. edulis* at a global scale (Borsa *et al.* 2007; Borsa *et al.* 2012), but were later shown to have been historically introgressed during secondary contact(s) caused by glacial oscillations (Roux *et al.* 2014).

### Difference of introgression rate in Chile caused by adaptive introgression?

Admixture profiles were remarkably stable across populations of allopatric and sympatric ranges. This widespread interspecific admixture suggest that range expansion of the two species, through both natural range shifts (with long-term environmental changes) and/or human-mediated introductions, occurred after a primary episode of contact between the two taxa, during which interspecific gene flow occurred. Genetic differentiations are however reported between allopatric and sympatric populations for the two species (Table 3) suggesting that intraspecific divergence history for each species influence more the genetic differentiation between populations than different rates of introgression between species. For example, the two sub-populations of North Sea (i.e. no.4a and 4b sampled both at Fiskebackskil at the surface and at 20m depth, respectively) exhibited a strong genetic differentiation with the other *C. intestinalis* populations and also between them (Fig.1b, Table S3) while they showed similar hybrid index values (Table 1). This strong genetic differentiation could be explained by a reduced gene flow between the two sub-populations in North Sea, a result which echoed to the pattern described in the doctoral thesis of Elin Renborg (https://gupea.ub.gu.se/handle/2077/35128). The poor connectivity is hypothesized to result of density discontinuity of sea water which separates shallow and deep populations of *C. intestinalis*. Such patterns of population differentiation have already been documented in other coastal marine species showing extended distribution along depth gradient (Jennings *et al.* 2013; Pivotto *et al.* 2015).

A noteworthy exception of the stability of admixture profiles is the *C. robusta* population from Chile which showed the highest number of loci with shared polymorphism with *C. intestinalis* (Fig.4a) and the highest *h*-values over all *C. robusta* populations (Table 1). Moreover, the position of the Chilean population on the first axis of the PCA (Fig.2a) first suggests residual genotypic covariance best explained by a higher level of introgression by *C. intestinalis* than other *C. robusta* populations. This is formally tested using TREEMIX and *f*_3_ statistical analyses (Fig.5, Table S5) which highlighted significant migration events between *C. intestinalis* ancestor and the Chilean population. Incomplete lineage sorting of ancestral polymorphism is not expected to create such asymmetry of shared polymorphism between populations, but point out evidence of local introgression in the Chilean population (Fraisse *et al.* 2016; Martin *et al.* 2013; Pickrell & Pritchard 2012). This pattern of local introgression is not uniformly distributed among loci (Figure S6), which is usually not accounted for in demographic inferences such as TREEMIX and could explain why the source of admixture is not a contemporary *C. intestinalis* population. This pattern could be a consequence of adaptive introgression in the genomic region of these introgressed loci, a process documented in several recent studies (Fontaine *et al.* 2015; Mendez *et al.* 2012; Pardo-Diaz *et al.* 2012). A similar pattern was observed in the *Mytilus* mussel complex of species where local introgression proved to be heterogeneous across loci (Fraisse *et al.* 2016). However, other processes can generate heterogeneous introgression rates such as heterogeneous load of deleterious mutations in migrant tracks (Christe *et al.* 2016; Harris & Nielsen 2016). None of the loci identified matched with genes coding for a known phenotypic or physiological trait (Table S4).

It is important to note that the first report of *C. robusta* along the Chilean coasts (with the name of *C. intestinalis* used until the recognition of *C. robusta* as a valid species) dates back to the middle of the 20^th^ century (Van Name 1945). We thus cannot exclude that local introgression have occurred in the source population(s) of the populations introduced in Chile rather than after the introduction (as an outcome of selection in the Chilean introduction range). A recent phylogeographic study based on mtDNA data (Bouchemousse *et al.* 2016a) pointed out a low genetic differentiation between populations of Chile and populations sampled in Japan, the putative native range of *C. robusta*. Further analyses are needed to investigate if this pattern could be due to adaptive introgression, using for instance modelling methods such as those performed by Fraisse *et al.* (2014) in a *Mytilus* sp. hybrid zone to examine the likelihood of adaptive introgression. A much larger number of population representatives of the global distribution of *C. robusta*, particularly populations of the Asian range, is also needed to investigate the processes that occurred in the SE Pacific as compared to the other regions where *C. robusta* is nowadays distributed.

In conclusion, our study confirmed the almost complete absence of contemporary gene flow in the human-mediated contact zone wherein *C. robusta* and *C. intestinalis* co-exist in sympatry/syntopy. Efficient reproductive barriers seem to prevent hybridization in the wild between the two species. These results are casting doubts that hybridization could impede the spread of the non-native. Ecological processes (e.g. niche displacement, trophic competition) might thus be more important to determine the fate of the two species in the sympatric range. Even if efficient reproductive isolation mechanisms are acting, few crosses involving an advantageous allele can be sufficient to favor its transmission in subsequent generations of the non-native species (Hedrick 2013). Our density of markers was clearly not sufficient to detect local signatures of adaptive introgression at genomic level. High-throughput genome analyses will be needed to definitively exclude, or confirm, that invasion potential of *C. robusta* is facilitated by adaptive introgression with *C. intestinalis* in the Northeast Atlantic. This approach might also allow us to identify genomic regions completely devoid of introgression, which may correspond to impassable reproductive barriers. Altogether, our study provides evidence that what was inferred to be recently introgressed individuals are more likely the outcome of a low level of residual historical introgression redistributed at global scale by natural range shifts and human-mediated introductions. Local introgression patterns, mostly concentrated on a few genome regions, were observed in the population sampled in the SE Pacific, a population far from the current distribution range of *C. intestinalis*. This result paves the way for further work to investigate adaptive introgression processes in other regions, in light of the range shift history of *C. robusta*.

## Acknowledgements

The authors are very grateful to C. Roux and N. Galtier for making available RNA sequences from the Pophyl Project, K. Belkhir of the Montpellier Bioinformatics Biodiversity computing platform for his precious help to the optimization of loci selection and the ADN^id^ society (Montpellier) for the genotyping of SNPs. We are also grateful to all our colleagues who contributed to the collection of samples: the divers of the Marine Operations department (*Service Mer & Observation*) at the Roscoff Biological Station, J.D.D. Bishop, S. Krueger-Hadfield, B. Lundve, J. Pechenik. The authors kindly acknowledge C. Roux and C. Fraisse for help and advices on R packages and scripts and three anonymous reviewers for their helpful comments and advices on the manuscript. This work benefitted from funding of the ANR project HYSEA (no. ANR-12-BSV7-0011) and the Interreg IVa Marinexus project, and a Languedoc-Roussillon Region “Chercheur(se)s d’avenir 2011” grant to NB.

## Data accessibility

Full dataset of the 310 SNPs were deposited into the DRYAD database (DOI: 10.5061/dryad.1h9b1).

## Author contributions

SB, CLH, NB and FV designed the study. CLH and NB with contribution of SB and FV performed the choice of SNP panel. SB and FV performed the choice of populations for genotyping. SB, CLH, NB and FV analyzed the data and wrote the article.

